# Do damaging variants of *SLC6A9*, the gene for the glycine transporter 1 (GlyT-1), protect against schizophrenia?

**DOI:** 10.1101/536904

**Authors:** David Curtis

## Abstract

**Aims:** To test whether genetic variants predicted to impair the functionality of *SLC6A9*, which codes for the GlyT-1 glycine transporter, are protective against schizophrenia.

**Method:** In an exome sequenced sample of 4225 schizophrenia cases and 5834 controls variants occurring in *SLC6A9* were annotated and weights were assigned using GENEVARASSOC. Genotype counts were compared using SCOREASSOC.

**Results:** Variants predicted to be deleterious by SIFT and damaging by PolyPhen were examined. Genotypes at 1:44466494-G/A seemed likely to be erroneous. If these were ignored then there were 15 damaging variants in controls and 5 in cases.

**Conclusions:** The results are consistent with the hypothesis that variants which damage *SLC6A9* are protective against schizophrenia but a larger sample would be required to confirm this.

**Declaration of interest:** The author declares no conflict of interest.

## Introduction

There is compelling evidence that impaired functioning of the glutamatergic N-methyl-D-aspartate receptor (NMDAR) can produce psychotic symptoms and is implicated in the pathogenesis of some cases of schizophrenia (Javitt and Zukin, 1991; Dalmau *et al.*, 2011; Steiner *et al.*, 2013; Schizophrenia Working Group of the Psychiatric Genomics Consortium, 2014; Genovese *et al.*, 2016; Curtis *et al.*, 2018; Tsavou and Curtis, 2019). The functioning of NMDAR can be enhanced by blockers of glycine transporter 1 (GlyT-1), which produce increased activation of the modulatory site at which glycine acts as a co-agonist (Hashimoto, 2014). GlyT-1 blockers have been trialled as treatments for schizophrenia. In a trial of bitopertine monotherapy compared against olanzapine or placebo in patients with an exacerbation of schizophrenia, all groups improved to a similar extent (Bugarski-Kirola *et al.*, 2014). Sarcosine, another GlyT-1 blocker, was superior to placebo in a number of studies (Tsai *et al.*, 2004; Lane *et al.*, 2005, 2010; Strzelecki, Urban-Kowalczyk and Wysokiński, 2018) but in others no difference was demonstrated (Lane *et al.*, 2006, 2008).

If blocking GlyT-1 has a therapeutic effect in schizophrenia then it would be reasonable to expect that genetic variants damaging *SLC6A9*, the gene which codes for it, might be protective against schizophrenia. Homozygous variants in *SLC6A9* can cause severe encephalopathy (Alfadhel *et al.*, 2016; Kurolap *et al.*, 2016) but no phenotypic effects have been described from heterozygous variants except for one report that rs16831558, which lies 2.1 kb upstream, may impact response to clozapine treatment (Taylor *et al.*, 2016).

The hypothesis that damaging variants of *SLC6A9* might be protective against schizophrenia was examined using data from a study of exome-sequenced cases and controls.

## Methods

The Swedish schizophrenia study, which had initially been tested for an excess of ultra-rare variants, was subsequently subjected to a weighted burden analysis, as reported previously (Genovese *et al.*, 2016; Curtis *et al.*, 2018). Whole exome sequence data was downloaded from dbGaP and, as explained previously, subjects with a substantial Finnish ancestry component were removed leaving a sample of 4225 cases and 5834 controls. Variants were annotated using VEP, PolyPhen and SIFT (Kumar, Henikoff and Ng, 2009; Adzhubei, Jordan and Sunyaev, 2013; McLaren *et al.*, 2016). After applying appropriate quality control measures, a weighted burden analysis of *SLC6A9* was performed using GENEVARASSOC and SCOREASSOC similar to the previous analysis, except that attention was not restricted to only rare variants (Curtis, 2012, 2016). GENEVARASSOC assigns functional weights to each variant based on the predicted effect on the functions of the gene so that, for example, a nonsynonymous variant is given a weight of 10 and a stop gained mutation a weight of 20. To these weights were added 10 if the variant was predicted to be deleterious by SIFT and 10 if it was predicted to be possibly or probably damaging by PolyPhen. SCOREASSOC then multiplies this functional weight by a frequency weighting factor so that very rare variants are weighted 10 times higher than common ones of the same type. The reasoning for including common variants in the analysis was that there might not be strong selection pressure against protective variants and hence they might not be particularly rare.

## Results

On examining the results of the weighted burden test, it was apparent that more of the variants with the highest weights were seen among controls rather than cases. There were no variants predicted to completely disrupt gene functioning, i.e. stop mutations or essential splice site variants, and so the variants with the highest weights were those nonsynonymous variants which were identified as both deleterious by SIFT and also as damaging by Polyphen. These variants are listed in Table 1 along with their genotype counts. VEP annotation was performed for each transcript of *SLC6A9* and the weights were assigned based on the most damaging annotation for any transcript. Table 1 provides an example of one of the transcripts with such an annotation.

**Table 1.** Genotype counts for variants in *SLC6A9* which are predicted both as deleterious by SIFT and as damaging by PolyPhen. For each variant an example transcript is shown which has both these annotations.

| Variant | Transcript | Effect | SIFT | Polyphen | ControlAA | ControlAB | ControlBB | CaseAA | CaseAB | CaseBB |
| --- | --- | --- | --- | --- | --- | --- | --- | --- | --- | --- |
| 1:44463225-G/A | ENST00000360584 | R705W | deleterious_low_confidence(0) | possibly_damaging(0.732) | 5802 | 1 | 0 | 4194 | 0 | 0 |
| 1:44463354-G/A | ENST00000360584 | R662W | deleterious(0) | possibly_damaging(0.863) | 5831 | 1 | 0 | 4223 | 0 | 0 |
| 1:44463544-C/T | ENST00000360584 | G637R | deleterious(0) | probably_damaging(0.943) | 5818 | 1 | 0 | 4199 | 0 | 0 |
| 1:44466481-G/T | ENST00000360584 | F571L | deleterious(0.02) | probably_damaging(0.998) | 5833 | 1 | 0 | 4224 | 0 | 0 |
| 1:44466494-G/A | ENST00000360584 | P567L | deleterious(0.04) | possibly_damaging(0.737) | 5816 | 18 | 0 | 4206 | 16 | 1 |
| 1:44466927-A/G | ENST00000360584 | V488A | deleterious(0.05) | possibly_damaging(0.669) | 5833 | 1 | 0 | 4225 | 0 | 0 |
| 1:44467115-G/T | ENST00000360584 | L456M | deleterious(0.04) | probably_damaging(1) | 5831 | 3 | 0 | 4222 | 1 | 0 |
| 1:44467139-G/A | ENST00000360584 | L448F | deleterious(0) | probably_damaging(1) | 5826 | 1 | 0 | 4220 | 0 | 0 |
| 1:44467157-C/T | ENST00000360584 | V442M | deleterious(0) | probably_damaging(0.989) | 5824 | 1 | 0 | 4214 | 0 | 0 |
| 1:44467202-C/T | XM_005271146.1 | V243L | deleterious(0.02) | \|probably_damaging(0.939) | 5814 | 1 | 0 | 4215 | 0 | 0 |
| 1:44467244-C/T | ENST00000360584 | V413I | deleterious(0) | probably_damaging(0.999) | 5826 | 0 | 0 | 4220 | 1 | 0 |
| 1:44467255-T/C | ENST00000360584 | Y409C | deleterious(0) | probably_damaging(1) | 5830 | 1 | 0 | 4222 | 0 | 0 |
| 1:44468249-C/T | ENST00000475075 | E154K | deleterious(0.04) | possibly_damaging(0.472) | 5833 | 1 | 0 | 4222 | 1 | 0 |
| 1:44475663-A/C | ENST00000360584 | V171G | deleterious(0) | possibly_damaging(0.811) | 5832 | 0 | 0 | 4223 | 1 | 0 |
| 1:44476409-C/T | ENST00000360584 | R132H | deleterious(0.01) | probably_damaging(1) | 5829 | 2 | 0 | 4223 | 0 | 0 |
| 1:44476469-G/A | ENST00000360584 | T112M | deleterious(0) | probably_damaging(1) | 5831 | 0 | 0 | 4221 | 1 | 0 |

Although if one counts the listed variants then more occur in controls than cases, it can be seen that 1:44466494-G/A is commoner than the others and occurs in 18 controls and 17 cases, one of which is homozygous for the alternate allele. Thus if this variant is included then the count of subjects possessing a variant allele does not differ much between controls and cases. However there are two reasons to question the validity of the genotypes for this variant. The first is that there is a subject who is called as a homozygote but one would only expect 3 subjects per million to be homozygous given that the observed frequency of the alternate allele is 0.0018. The second reason to be suspicious is that according to gnomAD the frequency for this allele among non-Finnish Europeans, based on 64084 subjects, is only 0.00039 and is even lower in other populations (Lek *et al.*, 2016). This is quite incompatible with the results we observe. Examination of the calls for this variant as provided in the VCF file does not reveal any obvious problems. All the calls are based on a large number of reads, with the minimum depth being 35 and roughly similar numbers of reads for each allele in the heterozygotes. The homozygote has 48 reads for the alternate allele and none for the reference. Nevertheless, given that the frequency is incompatible with ExAC and given that the observation of a homozygote would be extremely unlikely even with the observed frequency it seems at least plausible that the calls for this variant are simply wrong for some unknown reason.

If 1:44466494-G/A is ignored then variants predicted to severely affect functioning of *SLC6A9* are observed 15 times in controls and 5 times in cases, with the ratio of controls to cases being 5834/4225=1.4. Given that this is a *post hoc* observation it is not appropriate to provide a formal p value.

## Discussion

The results are consistent with the hypothesis that genetic variants severely affecting functioning of *SLC6A9* are protective against schizophrenia if one either assumes that the genotypes at 1:44466494-G/A are mistaken or that this variant does not actually have a severe effect in spite of the predictions of SIFT and PolyPhen. Predicting the effects of coding variants on gene functioning is an imprecise art and if one wanted more certainty about the impact of these variants one would need to study them in a model system such as cultured cells. The present small study is obviously not conclusive but can serve to generate predictions which can be tested as larger samples become available. If further evidence accrues that damage to *SLC6A9* is protective against schizophrenia then this might provide additional motivation to develop pharmacological interventions which target GlyT-1 or which seek to boost NMDAR functioning in other ways.

## Acknowledgements

The datasets used for the analysis described in this manuscript were obtained from dbGaP at http://www.ncbi.nlm.nih.gov/gap through dbGaP accession number phs000473.v2.p2. Samples used for data analysis were provided by the Swedish Cohort Collection supported by the NIMH grant R01MH077139, the Sylvan C. Herman Foundation, the Stanley Medical Research Institute and The Swedish Research Council (grants 2009-4959 and 2011-4659). Support for the exome sequencing was provided by the NIMH Grand Opportunity grant RCMH089905, the Sylvan C. Herman Foundation, a grant from the Stanley Medical Research Institute and multiple gifts to the Stanley Center for Psychiatric Research at the Broad Institute of MIT and Harvard.

